# Loss of Wt1 in the murine spinal cord alters interneuron composition and locomotion

**DOI:** 10.1101/226316

**Authors:** Danny Schnerwitzki, Sharn Perry, Anna Ivanova, Fabio Viegas Caixeta, Paul Cramer, Sven Günther, Kathrin Weber, Atieh Tafreshiha, Lore Becker, Ingrid L. Vargas Panesso, Thomas Klopstock, Martin Hrabe de Angelis, Manuela Schmidt, Klas Kullander, Christoph Englert

## Abstract

Rhythmic and patterned locomotion is driven by spinal cord neurons that form neuronal circuits, referred to as central pattern generators (CPGs). Recently, dI6 neurons were suggested to participate in the control of locomotion. The dI6 neurons can be subdivided into three populations, one of which expresses the *Wilms tumor suppressor gene Wt1*. However, the role that Wt1 exerts on these cells is not understood. Here, we aimed to identify behavioral changes and cellular alterations in the spinal cord associated with *Wt1* deletion. Locomotion analyses of mice with neuron-specific *Wt1* deletion revealed that these mice ran slower than controls with a decreased stride frequency and an increased stride length. These mice showed changes in their fore-hindlimb coordination, which were accompanied by a loss of contralateral projections in the spinal cord. Neonates with *Wt1* deletion displayed an increase in uncoordinated hindlimb movements and their motor neuron output was arrhythmic with a decreased frequency. The population size of dI6, V0 and V2a neurons in the developing spinal cord of conditional *Wt1* mutants was significantly altered. These results show that the development of particular dI6 neurons depends on *Wt1* expression and loss of *Wt1* is associated with alterations in locomotion.

## Introduction

In vertebrates, rhythmic activity is generated by a network of neurons, commonly referred to as central pattern generators (CPGs) (Grillner and Zangger 1979; Grillner 1985). CPGs do not require sensory input to produce rhythmical output; however, the latter is crucial for the refinement of CPG activity in response to external cues (Shik and Orlovsky 1976; Rossignol S 1988; Pearson 2003). The locomotor CPGs are located in the spinal cord and consist of distributed networks of interneurons and motor neurons (MN), which generate an organized motor rhythm during repetitive locomotor tasks like walking and swimming (Grillner 1985; McCrea and Rybak 2008).

The spinal cord develops from the caudal region of the neural tube. The interaction of secreted molecules including sonic hedgehog (Shh) and bone morphogenetic proteins (BMPs) provides instructive positional signals to the 12 progenitor cell domains that reside in the neuroepithelium (Alaynick et al., 2011). Each domain is characterized by the expression of specific transcription factor encoding genes that are used to selectively identify these populations. The dI1-dI5 interneurons are derived from dorsal progenitors and primarily contribute to sensory spinal pathways. The dI6, V0-V3 interneurons and MN arise from intermediate or ventral progenitors and are involved in the locomotor circuitry (Goulding 2009).

Whereas the involvement of V0 - V3 neurons in locomotion has been well documented, the role for dI6 neurons in locomotion has only recently been investigated (Andersson et al. 2012; Dyck et al. 2012). In particular, a part of the dI6 population shows rhythmically active neurons (Dyck et al. 2012), and a more defined subpopulation of dI6 neurons expressing the transcription factor *Dmrt3*, is critical for normal development of coordinated locomotion (Andersson et al. 2012). Another group of dI6 neurons is suggested to express the Wilms tumor suppressor gene *Wt1* but has not yet been characterized (Goulding 2009; Andersson et al. 2012).

*Wt1* encodes a zinc finger transcription factor that is inactivated in a subset of Wilms tumors, a pediatric kidney cancer (Call et al. 1990; Gessler et al. 1990). Wt1 fulfills a critical role in kidney development; however, the function of Wt1 is not limited to this organ. Phenotypic anomalies of *Wt1* knockout mice can be found, among others, in the gonads, heart, spleen, retina and the olfactory system (Kreidberg et al. 1993; Herzer et al. 1999; Moore et al. 1999; Wagner et al. 2002; Wagner et al. 2005). In one of the first reports on *Wt1* expression, the spinal cord was described as a prominent Wt1+ tissue (Armstrong et al. 1993; Rackley et al. 1993), however, until now there are no further reports regarding the function of Wt1 in the central nervous system (CNS).

Here, we have examined the role of Wt1 in the developing spinal cord. We performed locomotor analyses of conditional *Wt1* knockout mice and used molecular biological and electrophysiological approaches to elucidate the function of *Wt1* expressing neurons for locomotion. Our data suggest that *Wt1* expressing dI6 neurons contribute to the coordination of locomotion and that Wt1 is needed for proper dI6 neuron specification during development.

## Results

### Wt1 expressing cells in the spinal cord are dI6 neurons

In order to determine the spatial and temporal pattern of *Wt1* expressing cells in the spinal cord, we performed immunohistochemical analyses. Wt1+ cells were detected in the medioventral mantle zone of the developing spinal cord at embryonic day (E) 12.5 (Fig. 1A). Until E15.5, embryonic spinal cords showed a constant amount of Wt1+ cells; thereafter, their number gradually decreased until they could no longer be detected in adult mice (Fig. 1B).

**Figure 1.**
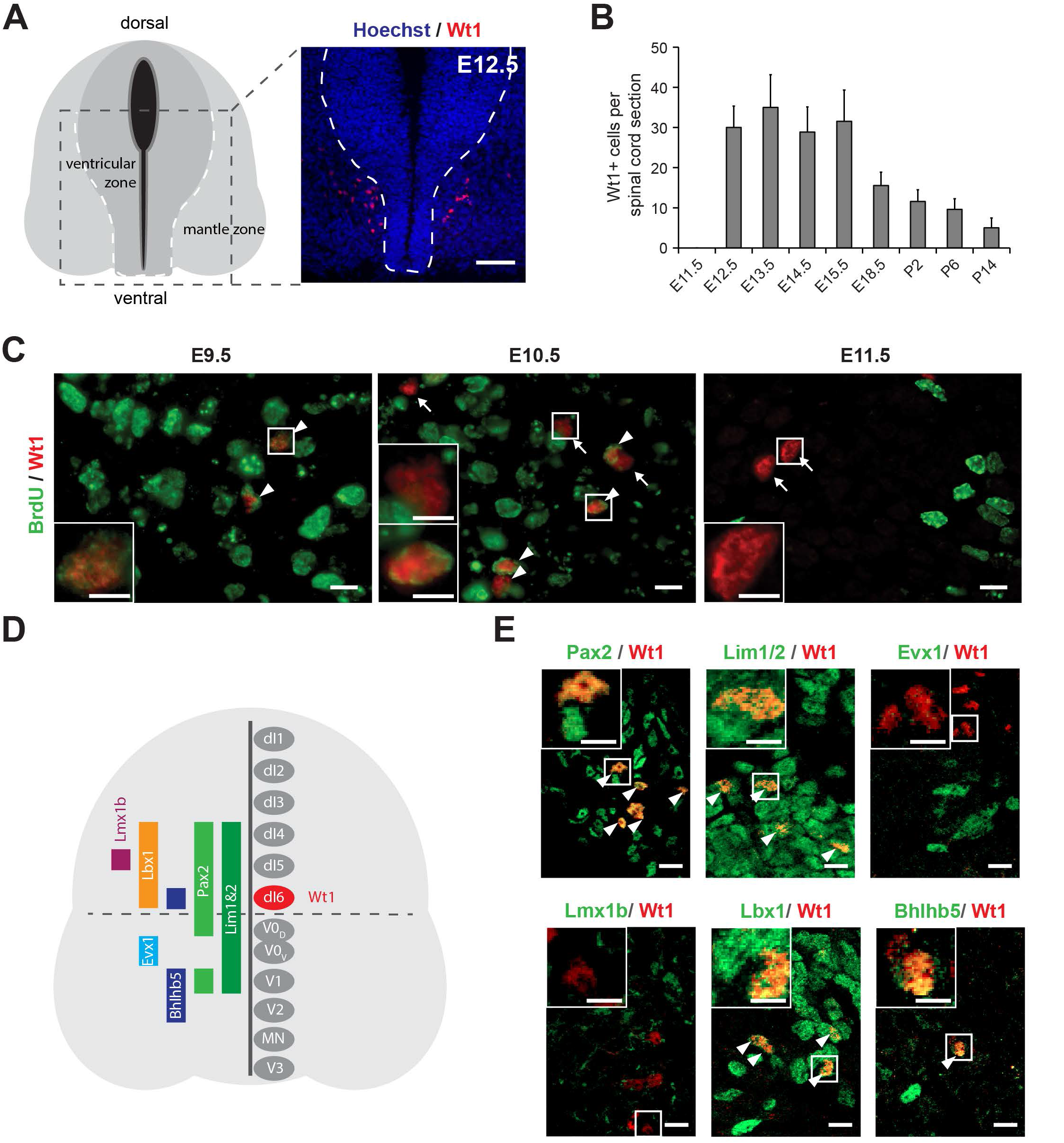
Characterization of Wt1+ neurons in the developing spinal cord. (A) Schematic illustration and Wt1 immunolabeling analysis of a transverse section (12μm) from E12.5 spinal cord showing the position of Wt1+ neurons (red) in the mantle zone of the developing spinal cord. Stippled line represents the border between the ventricular and mantle zones. Scale bar: 50 μm. (B) Plot showing the average cell number of Wt1+ neurons per 12 μm spinal cord section from different embryonic and postnatal stages. Wt1+ neurons are first found at E12.5 and decrease in cell number postnatally. (C) Determination of the birthdate of Wt1+ neurons by Bromodeoxyuridine (BrdU) proliferation assay. Proliferating cells situated in the ventricular zone were labelled by BrdU incorporation at different embryonic stages (E9.5, E10.5 and E11.5). Additional immunolabeling of these cells for Wt1 and BrdU at E12.5 revealed that prospective *Wt1* expressing cells still proliferate at E9.5 and at E10.5 but not at E11.5. Scale bar: 10 μm. Insets show higher magnifications of respective areas. Scale bar: 5 μm. (D) Schematic illustration of an E12.5 spinal cord section with markers and their occurrence in different neuron populations. These markers were used to establish the origin of Wt1+ neurons as dI6 neurons (red). (E) Immunolabeling of Wt1+ neurons with markers present in dI6 and adjacent interneuron populations. The partly overlapping location of Wt1 with Pax2, Lim1/2, Lbx1 and Bhlhb5 supports a dI6 character. Scale bar 10 μm. Insets show higher magnifications of respective areas. Scale bar: 5 μm.

We next wanted to determine the birthdate of Wt1+ cells, defined as the time point when progenitor cells cease to proliferate, leave the ventricular zone and start to differentiate. Using Bromodeoxyuridine (BrdU), the proliferative cells in the ventricular zone were labelled at different embryonic stages (E9.5, E10.5 and E11.5). Immunostaining of these cells for Wt1 at E12.5 revealed that prospective *Wt1* expressing cells still proliferate at E9.5 and even at E10.5 (Fig. 1C). At E11.5 Wt1 + cells no longer showed incorporation of BrdU suggesting that they had left the ventricular zone and started their migration and differentiation in the mantel zone at this time-point.

Wt1 has been proposed to label dI6 neurons (Goulding 2009), however, the only available primary data has so far only suggested its presence in a subpopulation of dI6 neurons expressing *Dmrt3* (Andersson et al., 2012). In order to closer examine the nature of Wt1+ cells, we performed immunostainings of embryonic spinal cords at E12.5. Cells expressing *Wt1* were positive for Pax2 and Lim1/2 labelling dI4, dI5, dI6, V0_D_ and V1 neurons (Tanabe and Jessell 1996; Burrill et al. 1997) while being negative for the post-mitotic V0_V_ marker Evx1 (Moran-Rivard et al. 2001) (Fig. 1D, E). *Wt1* expression did not overlap with Lmx1b, a marker specific for dI5 neurons, but did coincide with Lbx1 (Gross et al. 2002) and Bhlhb5 (Skaggs et al. 2011), which commonly occur in the ventral most dI4-dI6 Lbx1+ domain giving rise to dI6 neurons. Thus, these data supports and extends on the previous observations that Wt1 is a marker for a subset of dI6 neurons.

### Deletion of Wt1 affects locomotor behavior

To investigate the function of the Wt1+ neurons in the spinal cord, we made use of a *Nes-Cre;Wt1^fl/fl^* mouse line (Fig. 2A). At E12.5, no *Wt1* mRNA or protein was detected in neurons from this mouse line (Fig. 2A, B). Given the location of the Wt1 + neurons within the ventral dI6 population that has been shown to be involved in regulating locomotion, we performed behavioral tests associated with locomotion to investigate potential phenotypic consequences of deleting *Wt1* in spinal cord neurons. Footprints of adult mice walking on a transparent treadmill at fixed speeds (0.15, 0.25, 0.35 m/s) were recorded to analyze different gait parameters (Supplemental Fig. S1A). *Nes-Cre;Wt1^fl/fl^* mice revealed a significant reduction in stride frequency for both the fore- and hindlimbs relative to control (*Wt1^fl/fl^*) animals at all speeds measured. Heterozygous *Wt1* knockout mice (*Nes-Cre;Wt1^fl/^+*) did not differ significantly from controls. Stride length, accordingly, was significantly longer in *Nes-Cre;Wt1^fl/fl^* animals compared to wild type mice and *Nes-Cre;Wt1^fl/+^*. Thus, although *Nes-Cre;Wt1^fl/fl^* mice were slightly smaller compared to controls (body mass *Wt1^fl/fl^ vs* Nes-Cre;Wt1^fl/fl^ males 33 +/− 3.9 vs 25 +/− 3.7 g; females 25 +/− 3.2 g vs 22 +/− 1.4 g; body length males 9.9 +/− 0.4 g vs 9.4 +/− 0.4 cm; females 9.9 +/− 0.4 cm vs 9.8 +/− 0.3 cm), they made longer strides with lower frequency.

**Figure 2.**
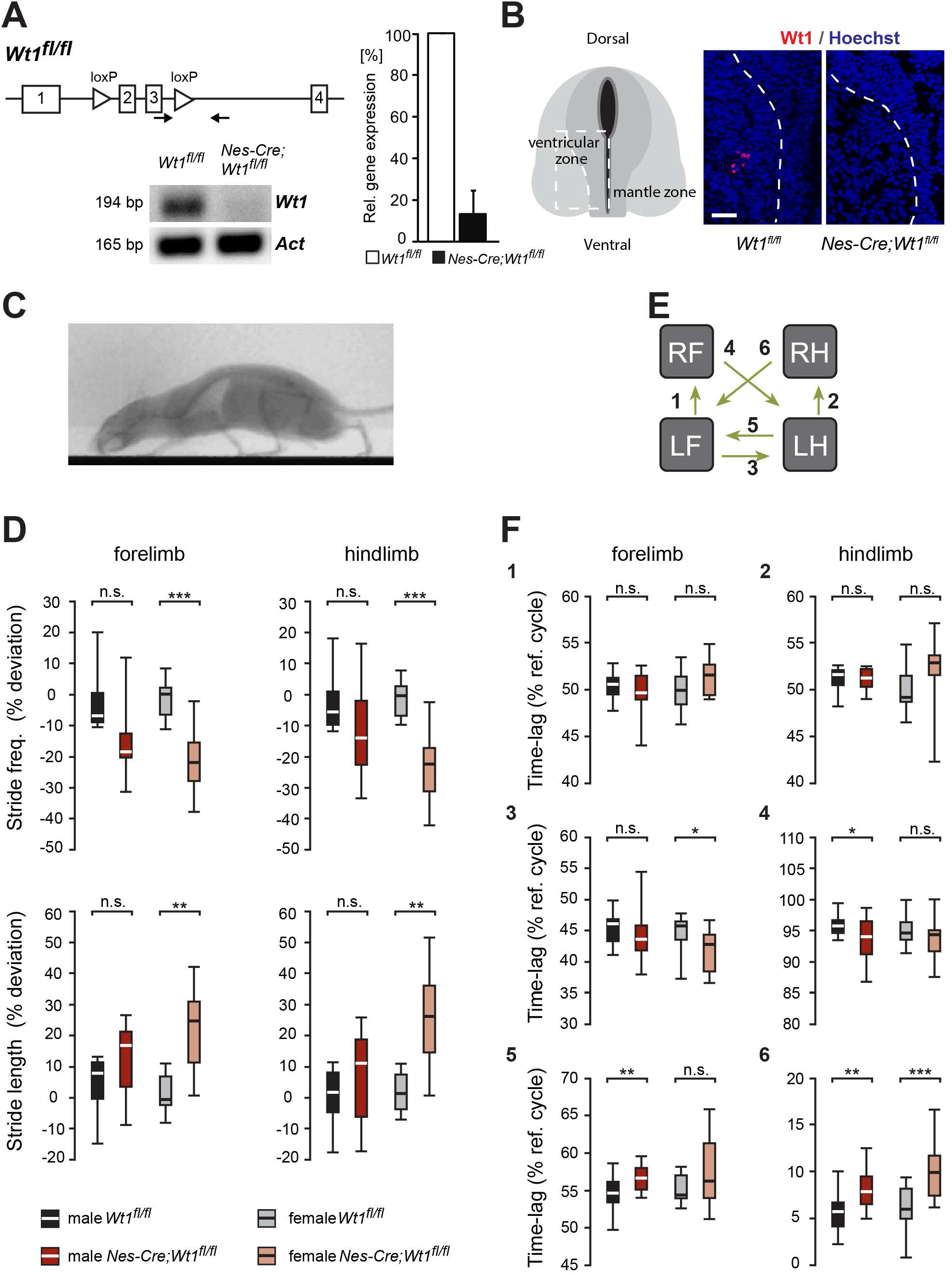
Mice with *Wt1* inactivation display altered locomotion. (A) Schematic illustration of the *Wt1^fl^* allele. *loxP* sites flanking exon 2 and 3 of the *Wt1* coding sequence allow Cremediated excision and conditional knockout of *Wt1*. Confirmation of a functional conditional *Wt1* knockout in *Nes-Cre;Wt1^fl/fl^* at E12.5 using qRT-PCR (quantification to the right). (B) Loss of Wt1 immunopositive signals in *Nes-Cre;Wt1^fl/fl^* embryos at E12.5 corroborates the loss of Wt1 protein. Schematic illustration shows the position where pictures were taken. Stippled line represents the border between ventricular and mantle zone. Scale bar: 40 μm. (C) X-ray radiograph of a walking mouse in lateral perspective. (D) Graphs displaying stride parameters collected in the X-ray radiograph. Stride frequency is significantly lower in female *Nes-Cre;Wt1^Ml^* mice in both forelimbs and hindlimbs. The stride length in female *Nes-Cre;Wt1^Ml^* mice is increased compared to female *Wt1^fl/fl^* mice, whereas smaller differences are found in male mice. Box plots indicate the median (bold white or black line), the 25^th^ and the 75^th^ percentile (box), and the data range (whiskers). Significance level of F_s_: *** P < 0.001; ** P < 0.01; * P < 0.05. (E,F) Interlimb coordination expressed as the time-lag between footfall events in percent stride duration. Left limbs are reference limbs. The scheme (E) illustrates, which phase relationships are shown by which graph. Phase relationships (F) between forelimbs (1) and between hindlimbs (2) illustrate overall symmetry of the walk. Timing of forelimb touchdown relative to hindlimb touchdown for ipsilateral (3) and contralateral (4) limbs show only minor differences between *Wt1^fl/fl^* and *Nes-Cre;Wt1^fl/fl^* mice. The timing of hindlimb footfalls relative to forelimb footfalls (5,6) differ between *Wt1^fl/fl^* and *Nes-Cre;Wt1^fl/fl^* mice, particularly at the contralateral limbs. Box plots indicate the median (bold white or black line), the 25^th^ and the 75^th^ percentile (box), and the data range (whiskers). Significance level: *** P < 0.001; ** P < 0.01; * P < 0.05.

To further explore gait alterations, we used X-ray fluoroscopy as a complementary method in a larger cohort of mice (Fig. 2C; Supplemental Fig. S1B; Supplemental Movie 1 and 2). When animals walked voluntarily at their preferred speed, deviations in stride frequency and stride length from the expected value (control baseline) for the given speed were again observed in *Nes-Cre;Wt1^fl/fl^* (Fig. 2D), but statistical significance is confirmed only for females. The changes were accompanied by a significant reduction of raw speed and size-corrected speed (= Froude number) in *Nes-Cre;Wt1^fl/fl^* mice of both sexes (Supplemental Fig. S1C). While both the duration of stance and swing phase and the distance covered by the trunk and the limbs, respectively, differ between controls and *Nes-Cre;Wt1^fl/fl^* by more than 10 percent in males and more than 15 percent in females, the ratio between the two phases, expressed by the Duty factor, remains unaffected (Supplemental Fig. S1D). Thus, the temporal coordination between stance and swing phase in adult *Nes-Cre;Wt1^fl/fl^* mice is normal.

We tested whether changes in gait parameters are accompanied by changes in the phase relationships between the limbs (Fig. 2E, F). The footfall pattern of control and *Nes-Cre;Wt1^fl/fl^* females did not show significant differences at the same speed of 0.21 m/s (Supplemental Fig 1E). However, the different spread along the X-axis indicates the evenly elongated stance and swing phases.

The symmetry of left and right limb movements expressed as the time-lag between footfalls in percent stride duration of a reference limb was unaffected in the *Nes-Cre;Wt1^fl/fl^* mice (Fig. 2E,F - 1 and 2). Also, the timing of forelimb footfalls relative to the hindlimb cycles is very similar between *Wt1^fl/fl^* mice and *Nes-Cre;Wt1^fl/fl^* mice (Fig. 2E,F – 3 and 4). Significant differences between *Wt1^fl/fl^* mice and *Nes-Cre;Wt1^fl/fl^* mice were observed in the timing of the hindlimb footfalls relative to the forelimb cycles (Fig. 2E,F – 3 and 4). The touchdown of the ipsilateral and the contralateral hindlimb fall in a later fraction of the forelimb stride cycle in *Nes-Cre;Wt1^fl/fl^* mice compared to the *Wt1^fl/fl^* mice. The deviation cannot be explained by the differences in animal speed, because the hind-to-forelimb coordination does not show speed-dependent variation.

**Figure 3.**
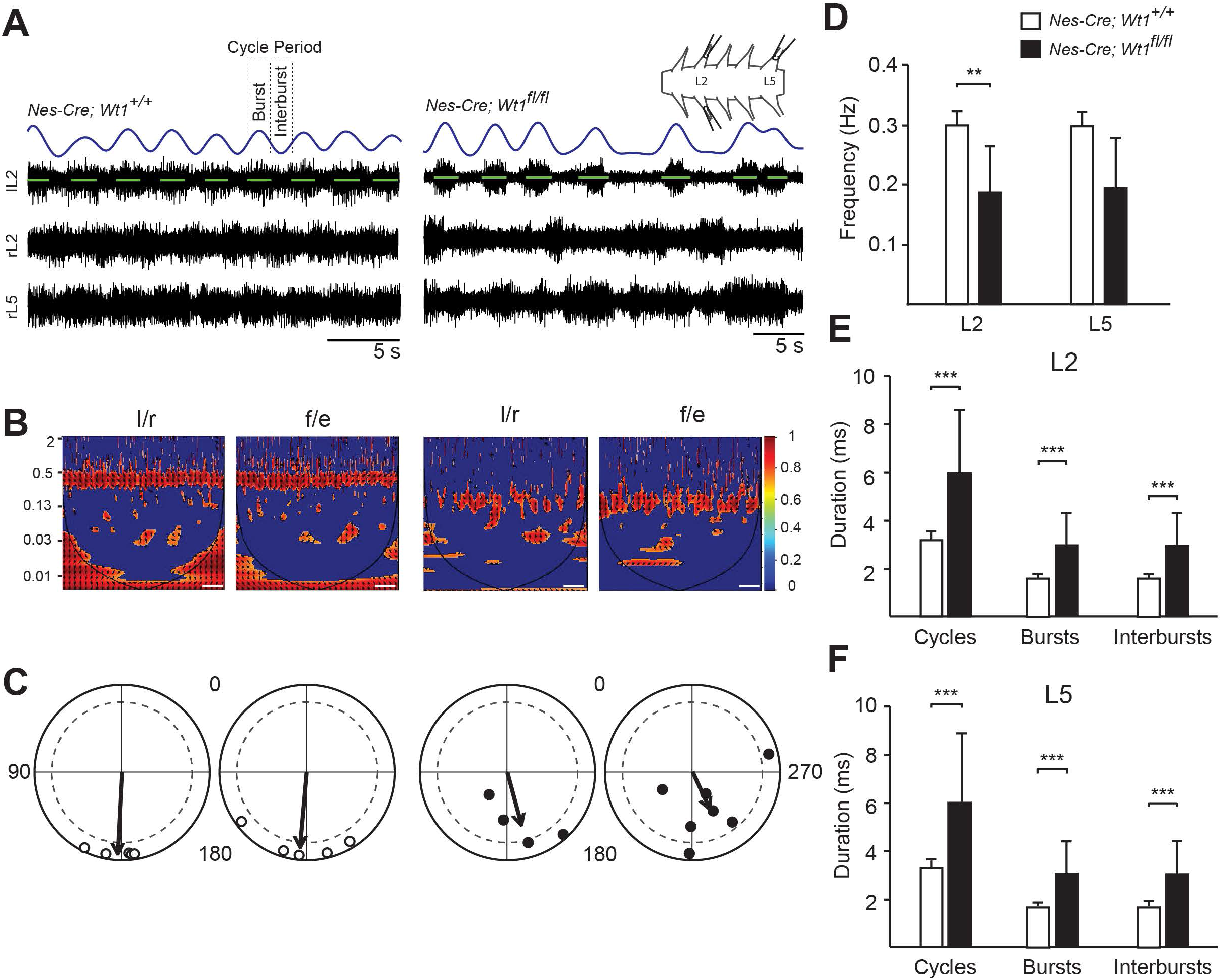
Locomotor activity is variable and uncoordinated in *Nes-Cre; Wt1^fl/fl^* pups. (A) Representative traces showing locomotor-like activity during fictive locomotion from left and right lumbar (L) 2 and right L5 ventral roots from control (*Nes-Cre; Wt1^+/+^*) and *Wt1* conditional knockout (*Nes-Cre; Wt1^fl/fl^*) mice. Rhythmic activity was induced by application of NMDA, serotonin and dopamine. Raw traces in black; rectified, low-pass filtered signal of lL2 trace in blue; activity burst shown in green. Spinal cord schematic depicts the attached suction electrodes to the right (r) and left (l) L2 and rL5 ventral roots. Scale bar: 5 seconds. (B) Phase analysis and associated coherence power spectra of left/right (L2/L2) and flexor/extensor (L2/L5) recordings. Regions of persistent coherence emerge for control mice at 0.30 Hz, whereas spinal cords from *Nes-Cre; Wt1^fl/fl^* mice show a reduced coherency region at 0.18 Hz. Color graded scale indicates normalized coherence. Scale bar: 125 seconds. (C) Locomotor patterns, analyzed from 20 consecutive bursts, reveal impaired and variable left/right and flexor/extensor alternation in *Nes-Cre; Wt1^fl/fl^* mice (black dots). Normal left/right and flexor/extensor alternation is maintained in control (white dots) mice. Each dot represents one cord; arrows represent the mean phase. The length of the vector is a measure of the statistical significance of the preferred phase; dashed grey line indicates region of high significance at 0.8 (Rayleigh test). Control, n = 5, *Nes-Cre;Wt1*^fl/fl^, n=7. (D) *Nes-Cre;Wt1^fl/fl^* mice have a slower locomotor frequency than control mice, p = 0.0060. (E – F) The slower locomotor frequency in *Nes-Cre; Wt1^fl/fl^* mice is mirrored by an increased cycle period, burst and interburst duration in both L2 (E) and L5 (F) roots. Data expressed as mean ± SD. Significance level: ** p < 0.01, *** p < 0.001.

**Figure 4.**
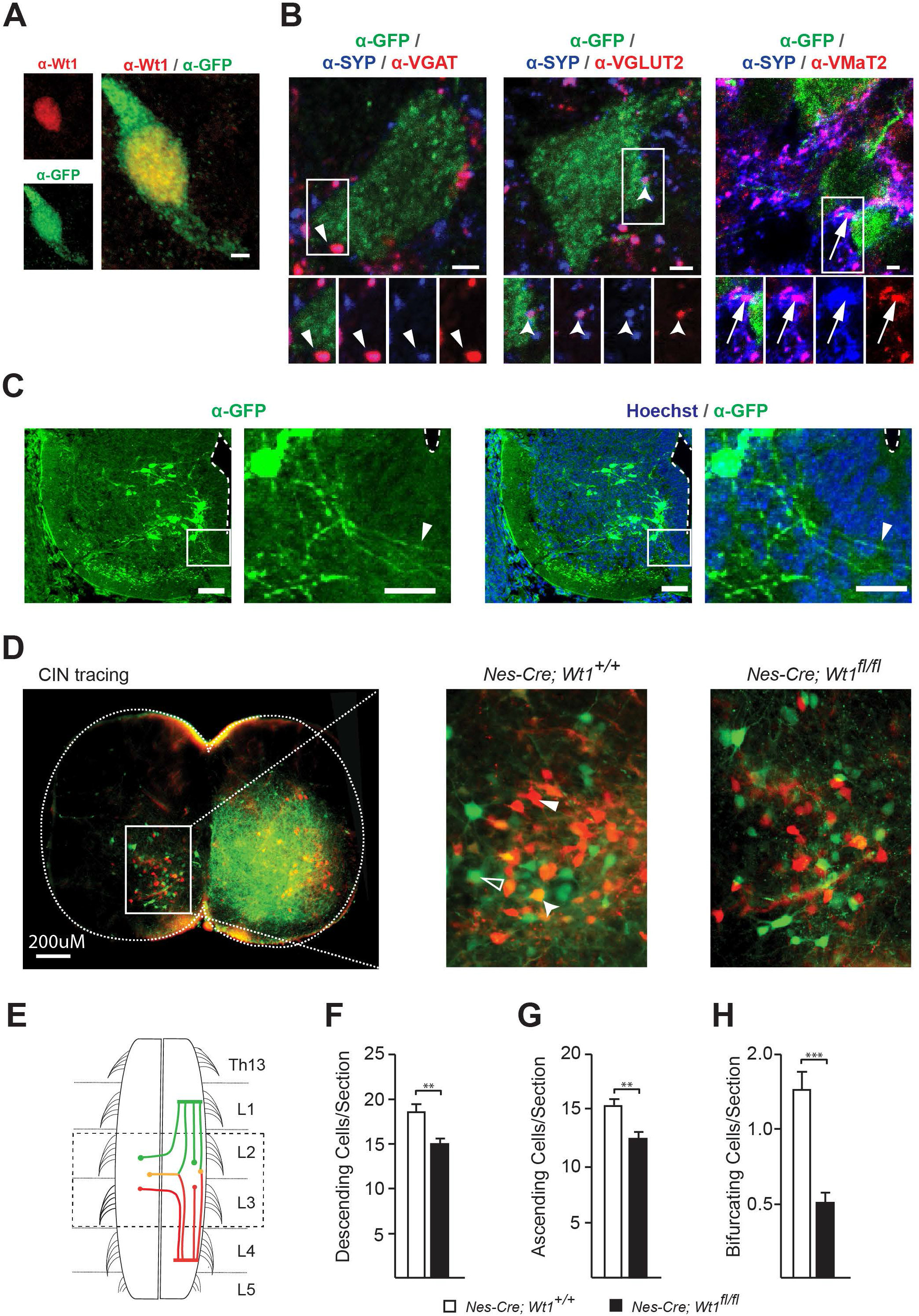
Innervation of Wt1+ neurons and number of commissural neurons in neonatal mice. (A) Spinal cords of *Wt1-GFP* embryos (stage E16.5) show a Wt1 + interneuron immunopositive for GFP. Wt1 is localized in the nucleus and GFP throughout the cell. Scale bar: 2 μm. (B) Wt1+ neurons (green) receive excitatory, inhibitory and monoaminergic synaptic contacts. Synaptic terminals are identified with synaptophysin (blue). Glutamatergic terminals were immunolabeled for VGLUT2; inhibitory synapses immunolabeled for VGAT, monoaminergic terminals immunolabeled for VMAT2. Arrows point to individual synaptic terminals (magenta) present on Wt1+ neurons (green). Boxed areas show higher magnification panels of separated channels. Scale bar: 2 μm. (C) GFP immunolabeled dI6 neurons in the spinal cord of E16.5 *Wt1-GFP* embryos. GFP antibody staining (green; left panel) and merge with Hoechst (blue; right panel) is shown. Boxed areas represent location of higher magnification panels shown on the right of each panel. Contralateral projections crossing the midline (dashed line) of the spinal cord are visible (arrow heads in magnified images). Scale bar: 20 μm for overview images and 50 μm for magnified images. (D) Photomicrographs of transverse, 60μm, lumbar, spinal cord sections with applied fluorescein-dextran amine (FDA, green) and Rhodamine-dextran amine (RDA, red) tracers. Higher magnification images of wild type (*Nes-Cre;Wt1^+/+^*) and homozygous (*Nes-Cre; Wt1^fl/fl^*) segments, showing intersegmental retrograde FDA (white arrow), RDA (open arrow) and double labelled (triangle arrow) neurons. Scale bar: 200μm. (E) Schematic illustration of FDA (lumbar (L)1) and RDA (L4/5) application sites tracing descending (green), ascending (red) and bifurcating (yellow) neurons. The area of analysis (L2/3) is indicated by black dashed line. (F – H) Quantification of descending FDA labelled neurons (F), ascending RDA labelled neurons (G), and bifurcating, double labelled neurons per section (H). Descending, ascending and bifurcating CINs are significantly fewer in homozygous spinal cords compared to control cords. Data expressed as mean ± SEM. Significance level:*P < 0.05, **P < 0.01, ***P < 0.001.

So far, the limb kinematics of adult *Nes-Cre;Wt1^fl/fl^* mice compared to the *Wt1^fl/fl^* mice shows subtle differences in gait parameters and interlimb coordination with a high degree of variation. In sum, these differences result in a performance reduction indicated by the overall lower walking velocities.

### Deletion of Wt1 results in a disturbed and irregular postnatal locomotor pattern

After having observed altered gait parameters in adult *Nes-Cre;Wt1^fl/fl^* animals, we wondered whether gait also would be affected in younger mice. Indeed, *Nes-Cre;Wt1^fl/fl^* pups had more difficulty coordinating their fore- and hindlimbs compared to controls when performing air stepping. Although there was no increase in hindlimb synchronous steps, left/right alternating steps were decreased and the number of uncoordinated steps were increased in *Nes-Cre;Wt1^fl/fl^* animals (Supplemental Fig 2; Supplemental Movie 3 and 4). We next performed fictive locomotion experiments on isolated spinal cords from control and *Nes-Cre;Wt1^Ml^* mice (P0-P3). Fictive locomotor drugs induced a markedly slower, disturbed, more variable pattern of locomotor-like activity in *Nes-Cre;Wt1^fl/fl^* spinal cords (n = 6) compared to the stable, rhythmic pattern of locomotor-like activity in control mice (n = 5). Control spinal cords had recorded activity bursts that showed clear left/right (L2 vs L2) and flexor/extensor (L2 vs L5) alternation that persisted throughout activity periods, whereas activity bursts in *Nes-Cre;Wt1^fl/fl^* spinal cords were uncoordinated and did not maintain strict left/right or flexor/extensor alternation (Fig 3A-B). The relationship between left/right and flexor/extensor alternation was examined and gave a strong phase preference for alternating bursts in control (Fig. 3C; l/r control, average phase preference: 183.4° R = 0.93; f/e control, 185.2°, R = 0.84). However, spinal cords from mice with a *Wt1* deletion showed an irregular locomotor pattern with inconsistent alternation as indicated by the length and direction of the phase vector (Fig. 3C; l/r average phase preference: 165.3°, R = 0.60; f/e 155.2°, R = 0.44). Additionally, the frequency of the ventral root output was decreased (Fig 3D: control; 0.30 ± 0.024 Hz: *Nes-Cre;Wt1^fl/fl^;* 0.18 ± 0.08 Hz). This slower rhythm in *Nes-Cre;Wt1^fl/fl^* cords could be attributed to altered L2 and L5 activity burst parameters, as *Nes-Cre;Wt1^fl/fl^* mice had significantly longer burst, interburst and cycle periods compared to control (Fig. 3E, F). Thus, the deletion of Wt1 results in a disturbed and irregular locomotor pattern, which suggests that there are changes to the neuronal locomotor circuitry that occur following *Wt1* deletion.

### Wt1+ neurons receive various synaptic inputs and can project commissurally

In order to assess how Wt1+ dI6 neurons are connected within the CPG network, we focused on the innervation pattern of these cells. We used the *Wt1-GFP* reporter mouse line (Hosen et al. 2007) where Wt1+ neurons are labeled by GFP. In contrast to the restricted localization of Wt1 in the nucleus, GFP is distributed throughout the cytoplasm and labels the soma and major processes (Fig 4A). In combination with antibodies against particular vesicular synaptic transporters, we observed that excitatory (VGLUT2), inhibitory (VGAT) and modulatory (VMaT2) synapses contact the soma of Wt1+ dI6 neurons (Fig. 4B). This shows that Wt1+ dI6 neurons receive excitatory, inhibitory and modulatory inputs suggesting that Wt1 + neurons are positioned to receive a multitude of signals and could act during locomotion to integrate different CPG signals.

Using the *Wt1-GFP* reporter mouse, we found GFP+ fibers crossing the spinal cord midline beneath the central canal suggesting that Wt1+ neurons project commissural fibers (Fig. 4C). Fluorescent dextran amine retrograde tracing of contralateral projections confirmed that at least part of the Wt1+ dI6 neurons project commissurally (Supplemental Fig 3). We analyzed spinal cord commissural neurons in control (*Nes-Cre;Wt1^+/+^*) and homozygous (*Nes-Cre;Wt1^fl/fl^*) mice (P1-5) to determine if the deletion of *Wt1* alters the total number of commissural neurons and investigated ascending (aCIN), descending (dCIN) and bifurcating (adCIN) subpopulations (Fig. 4 D, E). All traced subpopulations were markedly reduced in *Nes-Cre;Wt1^fl/fl^* cords spinal cords compared to controls (Fig. 4F – H).

### Loss of Wt1 leads to altered interneuron composition

To assess the possible impact of *Wt1* deletion for interneuron development, we analyzed dI6 and non-dI6 populations situated in the embryonic ventral spinal cord. The number of *Dmrt3* expressing cells, which constitutes a distinct but partly overlapping dI6 population (Andersson et al. 2012), was significantly decreased in the embryos harboring a loss of Wt1 in the spinal cord already at E12.5 (Fig. 5A) persisting throughout development (E16.5 and P1). At any investigated time point, neurons co-expressing both Wt1 and Dmrt3 were not detected in *Nes-Cre;Wt1^fl/fl^* embryos and neonates.

**Figure 5.**
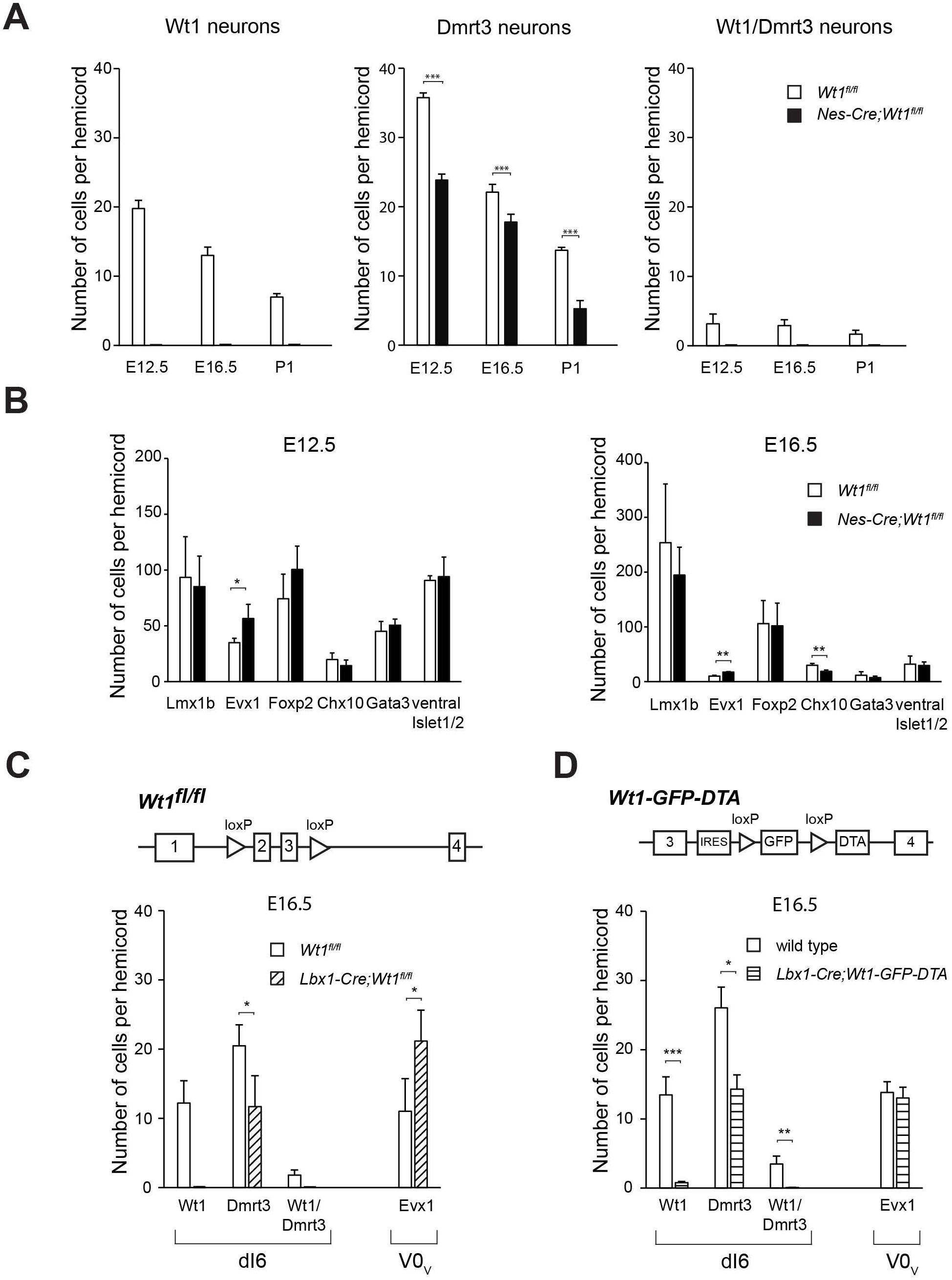
Alterations in the composition of ventral neurons upon *Wt1* knockout. (A) Average cell number of Wt1-, Dmrt3- and Wt1/Dmrt3 neurons per 12 μm spinal cord section from different embryonic and postnatal stages of control (*Wt1™*) and *Wt1* conditional knockout (*Nes-Cre;Wt1^fl/fl^*) mice. Number of Dmrt3 neurons significantly decrease in *Nes-Cre;Wt1^fl/fl^*. No Wt1/Dmrt3 neurons are detected in *Nes-Cre;Wt1^fl/fl^* animals. (B) Average cell number of Lmx1b, Evx1, Foxp2, Chx10, Gata3 and ventral Islet1/2 neurons per 12 μm spinal cord section from control (*Wt1^fl/fl^*) and homozygous (*Nes-Cre;Wt1^fl/fl^*) mice at E12.5 and E16.5. Number of Evx1+ V0 neurons is significantly increased. (C) Average cell number of Wt1-, Dmrt3- and Wt1/Dmrt3 dI6 neurons and Evx1 V0 neurons per 12 μm spinal cord section from E16.5 control (*Wt1^fl/fl^*) and *Wt1* conditional knockouts (*Lbx1-Cre;Wt1^fl/fl^*) mice. *Lbx1-Cre* based conditional *Wt1* knockouts show a decrease in the amount of dI6 neurons and an increase in the cell number of Evx1 neurons as for *Nes-Cre; Wt1^fl/fl^* animals. (D) Schematic illustration of *Wt1-GFP-DTA* allele. Cassette consisting of *loxP* sites flanking *GFP* coding sequence upstream of *Diphtheria toxin subunit A (DTA*) was inserted into the *Wt1* locus. This cassette allows Cre-mediated ablation of Wt1 + neurons. Graph shows average cell number of Wt1-, Dmrt3- and Wt1/Dmrt3 dI6 neurons and Evx1+ V0 neurons per 12 μm spinal cord section from E16.5 wild type control and *Lbx1-Cre; Wt1-GFP-DTA* mice. Nearly all Wt1+ neurons are absent. The number of Dmrt3 neurons is significantly decreased. Population size of Evx1 neurons is not altered after ablation of Wt1+ neurons. Data expressed as mean ± SD. Significance level: *P < 0.05, **P < 0.01, ***P < 0.001.

Loss of the transcription factor Dbx1 that is involved in differentiation of the V0 population results in a fate switch of some V0 neurons to become dI6 interneuron-like cells (Lanuza et al., 2004). Thus, we investigated whether populations flanking the dI6 population were affected in *Nes-Cre;Wt1^fl/fl^* mice. The Lmx1b+ dI5 population was similar in number when comparing *Nes-Cre;Wt1^fl/fl^* with wild type embryos, whereas the number of Evx1+ V0_V_ neurons was significantly increased already at E12.5 (Fig. 5B). This increase was still detectable at E16.5. No differences could be seen in Foxp2+ V1 neurons, Chx10 (V2a) and Gata3 (V2b) neurons and Islet 1/2+ motor neurons between conditional *Wt1* knockout and control embryos at E12.5. However, at E16.5 Chx10+ V2a neurons showed a significant decrease in cell number.

To verify the changes of interneuron composition found in the developing *Nes-Cre;Wt1^fl/fl^* mice, we made use of a second mouse line, namely *Lbx1-Cre;Wt1^fl/fl^* mice. At embryonic stage E16.5, we observed a decrease in the amount of dI6 neurons and an increase in the cell number of Evx1+ neurons similar to *Nes-Cre;Wt1^fl/fl^* mice (Fig. 5C). This decline in the number of dI6 neurons and the concomitant increase in the amount of Evx1+ neurons might point to a change in the developmental fate from dI6 neurons into V0 neurons prompted by deletion of *Wt1*. To test this hypothesis, we ablated the cells destined to express *Wt1*. We used *Lbx1-Cre;Wt1-GFP-DTA* mice in which the Diphtheria toxin subunit A (*DTA*) is expressed from the endogenous *Wt1* locus after Cre-mediated excision of a *GFP* cassette harboring a translational STOP-codon. *Cre* expression driven by the *Lbx1* promoter targets the dI4 to dI6 interneuron populations (Müller et al. 2002). In *Lbx1-Cre;Wt1-GFP-DTA* embryos, nearly all Wt1+ neurons were ablated at E16.5 (Fig. 5D). The ablation of Wt1+ neurons coincided with a significantly decreased number of Dmrt3+ neurons in *Lbx1-Cre;Wt1-GFP-DTA* embryos, but did not affect the number of Evx1 + neurons (Fig 5D). Taken together, the results from the *Wt1* deletion and the ablation of the Wt1 neurons suggests that the fate switch from dI6 neurons into Evx1+ V0 neurons occurs due to the deletion of *Wt1*. A postnatal phenotypic behavioral analysis of these mice was not possible because neonates died immediately after birth due to serious respiratory deficits (data not shown).

The analyses of the interneuron composition in developing conditional *Wt1* knockout mice and embryos with an ablation of Wt1+ neurons suggest a fate switch within a specific subset of dI6 and V0_V_ neurons that depends on the presence of the cells destined to express *Wt1*.

### The transition of dI6 neurons into Evx1+ V0_V_ neurons upon loss of Wt1 is not direct

In order to further investigate the cellular fate change upon deletion of *Wt1* we combined *Wt1-GFP* and *Nes-Cre;Wt1^fl/fl^* animals to generate *Nes-Cre;Wt1^fl/GFP^* mice. These mice harbor a constitutive knockout allele of *Wt1* due to the insertion of a *GFP* coding sequence and another conditional *Wt1* knockout allele. GFP and Wt1 were co-localized in the ventral spinal cord of *Wt1^fl/GFP^* control animals at E13.5, whereas GFP, but not Wt1, was detected in spinal cords of *Nes-Cre;Wt1^fl/GFP^* embryos of the same age (Fig 6A). Thus, *Nes-Cre;Wt1^fl/GFP^* mice allowed us to inactivate *Wt1* while the cells destined to express *Wt1* are labelled by GFP.

**Figure 6.**
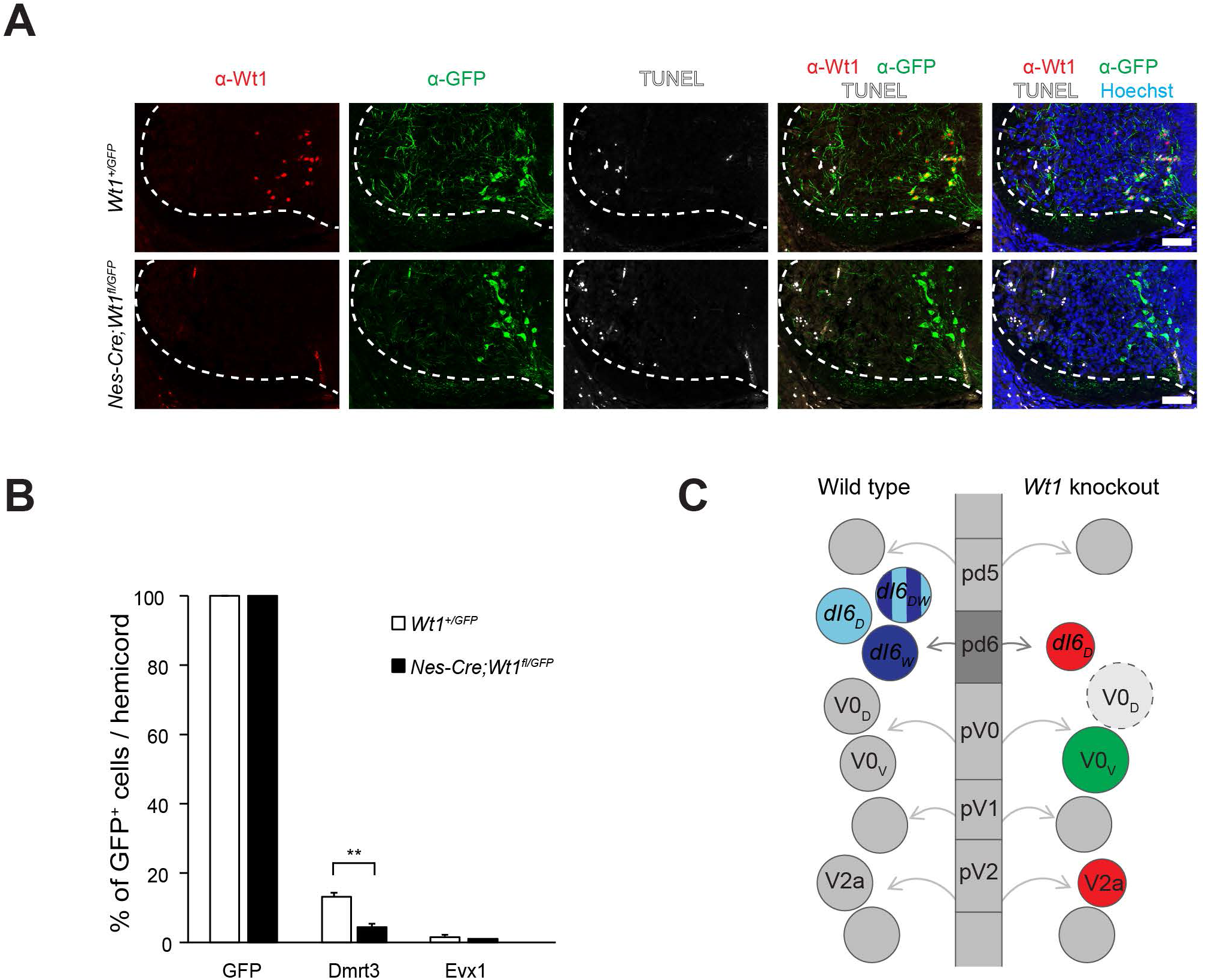
(A) Immunofluorescence staining of spinal cord sections from E13.5 *Wt1+^/GFP^* (control) and *Nes-cre;Wt1^fl/GFP^* embryos. GFP is depicted in green, Wt1 in red, TUNEL+ cells in white and Hoechst in blue. Orientation: dorsal at the top and ventral at the bottom. Scale bar: 50 μm. (B) Quantification of GFP+ cells harbouring the interneuron markers Dmrt3 and Evx1. Analysis was performed using E12.5 *Wt1+^/GFP^* (control; n=3) and *Nes-cre;Wt1^fl/GFP^* (n=3) embryos. The number of cells showing co-localization of GFP and the respective markers was determined and normalized to the total number of GFP+ cells, which was set to 100%. Upon *Wt1* knockout the amount of GFP+ cells possessing Dmrt3 is significantly decreased. The amount of GFP+ cells possessing Evx1 is not altered, suggesting an indirect fate change of dI6 neurons into Evx1+ V0_V_-like neurons upon *Wt1* knockout. Data expressed as mean ± SD. Significance level: **P < 0.01. (C) Scheme represents various progenitor cell domains (pd5, pd6, pV0, pV1 and pV2) that give rise to different populations of spinal cord neurons (shown as circles) in wild type and tissue specific *Wt1* knockout. In wild type animals, progenitor cells leave these domains, become postmitotic and differentiate into distinct interneuron populations that further subdivide. The dI6 interneuron population consists of neurons either positive for Dmrt3 (dI6_D_), Wt1 (dI6_W_) or both (dI6_DW_). Due to the knockout of *Wt1*, no dI6_W_ and dI6_DW_ are detectable and the number of dI6_D_ neurons is decreased. In contrast, the number of Evx1+ V0_V_ neurons increases, which is an indirect effect as potential dI6_W_ cells that lack Wt1 did not show a Evx1 signal. This effect might be explained by a hypothetical fate change of dI6 neurons into V0_D_ like neurons (dashed light grey circle). The increased number of V0_D_ neurons would thus prompt the pV0 progenitor cells to differentiate preferentially into V0_V_ neurons, which would compensate the excess amount of V0D neurons and lead to an increase in the population size of Evx1+ V0_v_ neurons. As a secondary effect, the number of V2a neurons, which innervate the V0_V_ neurons, declines at later developmental stages when neurons start to connect to each other potentially compensating the increased number of V0_V_ neurons. Only the subsets of interneuron populations are shown that are affected by the tissue specific *Wt1* knockout. Red indicates decrease in population size, green indicates increase in population size.

To investigate whether *Wt1* deletion leads to apoptosis in the respective cells, terminal deoxynucleotidyl transferase dUTP nick end labeling (TUNEL) was used. TUNEL+ cells were present in the ventrolateral spinal cords of *Wt1^fl/GFP^* control and *Nes-Cre;Wt1^fl/GFP^* embryos (Fig 6A). However, TUNEL signals never overlapped with GFP+ dI6 neurons destined to express *Wt1*, suggesting that *Wt1* inactivation in dI6 neurons did not result in cell death.

In order to find out whether cells destined to express *Wt1* would directly convert to V0_V_ neurons upon Wt1 inactivation, we performed immunohistochemical analyses. The presence of Dmrt3 and Evx1 in GFP+ dI6 neurons was analyzed in *Wt1^fl/GFP^* control and *Nes-Cre;Wt1^fl/GFP^* embryos at E12.5 (Fig 6B). The number of GFP+ cells per hemicord was determined and set to 100%. The proportion of Dmrt3+ cells was approximately 13% of all GFP+ cells in spinal cord of E12.5 control embryos. When *Wt1* was absent, the amount of Dmrt3+ GFP cells significantly decreased to 4%. In contrast, the proportion of GFP+ dI6 neurons that also showed Evx1 staining was not changed between *Wt1^fl/GFP^* control and *Nes-Cre;Wt1^fl/GFP^* animals (below 1% for both). Thus, the increase in the amount of Evx1+ V0_V_ neurons observed in mice lacking Wt1, does not seem to result from a direct transition of future Wt1+ dI6 neurons into Evx1+ V0_V_ neurons.

## Discussion

In this study, we have examined Wt1, which marks a subset of dI6 neurons. We found that Wt1 is required for proper differentiation of spinal cord neurons and that deletion of *Wt1* results in locomotor aberrancies in neonate and adult mice.

Neonates lacking Wt1 in the spinal cord increased the number of uncoordinated steps, which was supported by a markedly slower and variable pattern of locomotor-like activity in isolated *Nes-Cre;Wt1^fl/fl^* spinal cords. Adult *Nes-Cre;Wt1^fl/fl^* animals showed an increased stride length and a decreased stride frequency resulting in slower absolute locomotor speed, however, these alterations were more subtle compared to locomotion abnormalities seen in neonates. Compensatory adaptation of functional properties during postnatal maturation of neuronal circuits could possibly act to reduce the severity of the deficits seen in neonates. For instance, the corticospinal tract, which allows control of the spinal circuits directly by the motor cortex and overrides the spinal circuits, does not reach the lumbar spinal cord until approximately P7-P9 (Kamiyama et al. 2015).

Wt1+ neurons are unlikely to participate in locomotor rhythm generation *per se* since a rhythm is established when *Wt1* is deleted. We hypothesize that these neurons are involved in the maintenance or modulation of this rhythm. Adult *Nes-Cre;Wt1^fl/fl^* animals show a decreased stride frequency. The hindlimb-to-forelimb phase relationship is altered in animals with *Wt1* deletion in the spinal cord. This supports a possible role of the Wt1+ dI6 neurons in both timing and limitation of the stride cycle. An involvement in timing of the stride cycle would require an integrative position in the locomotor CPGs, which is compatible with the observed multi-synaptic input to Wt1+ dI6 neurons.

The timing of hindlimb footfalls relative to forelimb footfalls differed between *Wt1^fl/f^* and *Nes-Cre;Wt1^fl/fl^* mice, particularly at the contralateral limbs, suggesting Wt1 cells to play a role for long range coordination between various spinal cord segments. This phenotype is supported by the observation that at least a fraction of Wt1+ dI6 neurons possess commissural projections and thus are involved in the contralateral communication between the spinal cord halves. If and how the loss of this communication affects the adult locomotor phenotype does not become apparent from our data set of footfall timing but we expect to detect more details by the analysis of intrinsic limb kinematics. Deletion of *Wt1* leads to a decline in the number of commissural neurons suggesting an involvement of Wt1 in establishing proper projections of the *Wt1+* dI6 neurons. Thus, it will be of future interest to investigate the transcriptome of Wt1+ dI6 neurons and screen for potential target genes involved in axon guidance.

Lack of *Wt1* in the spinal cord causes alterations in the differentiation of dI6, V0 and V2a spinal cord neurons (Fig 6C). The inverse alterations in the dI6 and V0 populations suggest a fate change from dI6 to V0-like neurons when *Wt1* is inactivated. The putative transition from dI6 to V0-like neurons occurs at the time-point when *Wt1* expression would normally start. This instantaneous effect might be due to the derivation of both interneuron populations from neighboring progenitor domains sharing common transcription factors such as Dbx2 (Alaynick *et al*., 2011). Thus, loss of *Wt1* might lead to a switch in developmental programs that are normally repressed; whether this repression occurs cell-autonomously or non-cell-autonomously still has to be determined. In any case, when future *Wt1+* cells are ablated an increase of V0-like neurons is no longer observed, suggesting that the fate switch requires the cells about to express *Wt1*.

The fate change of prospective dI6 to V0-like neurons is complex since dI6 neurons can be subdivided into at least three subsets based on expression of the transcription factor encoding genes *Wt1* and *Dmrt3* (Fig 6C). Loss of *Wt1* not only affects the small number of dI6 neurons that possess Wt1 and Dmrt3 but also the number of neurons that only express *Dmrt3*. This population is significantly decreased. The presence of Wt1+ neurons therefore is essential to maintain the character of a subset of Dmrt3 neurons. If *Wt1* is inactivated, in addition to the cells which are programmed to express *Wt1*, possibly also Dmrt3+ neurons may differentiate into V0-like neurons.

Two main subpopulations exist within the V0 population (Alaynick *et al*., 2011): the Evx1+, more ventrally derived V0_V_ and the Evx1 negative, more dorsally derived V0D population, for which no distinct marker has yet been described. The knockout of *Dbx1* results in loss of the whole V0 population, whereby Evx1+ V0_V_ neurons acquire a more ventral fate and become V1 neurons, whereas Evx1 negative V0_D_ neurons acquire characteristics of dI6 neurons (Lanuza *et al*., 2004). This suggests that the V0_D_, rather than the V0_v_, neurons closer resemble the dI6 neurons and poses the question whether the fate change from Wt1+ dI6 neurons to Evx1+ V0_V_-like neurons represents a direct or an indirect transition. The investigations using the *Nes-Cre;Wt1^fl/GFP^* mice suggest that the Wt1-deficient dI6 cells do not change their fate directly into Evx1+ V0_V_-like neurons suggesting an indirect transition. That points to the possibility that the fate change might be achieved by a transition of Wt1+ dI6 neurons into the more closely related Evx1 negative V0_D_-like neurons, which leads to a putative increase of the V0_D_ population (Fig 6C). The Evx1+ V0_V_ population might, in turn, increase its number to compensate for a higher proportion of V0_D_-like neurons.

In addition to the changes in the dI6 and V0 population that occur upon *Wt1* deletion in the spinal cord, Chx10+ V2a neurons show a slight but significant decrease in their cell number at E16.5 (Fig 6C). This might represent a secondary effect of the alterations in the dI6 and V0 population, which occur already at E12.5. It was reported that V2a neurons directly innervate V0_V_ neurons (Crone *et al*., 2008). This secondary effect might thus be due to a potential adaptation to the altered interneuron composition in the spinal cord and the necessity to form proper contacts with target cells to build up the neuronal circuits responsible for locomotion.

In sum, the results obtained in this study not only shed light on the so far undescribed necessity for Wt1 in the development of spinal cord neurons and their functional implementation in circuits responsible for locomotion. The data also broadens our view on the complex interplay of the various neuron subpopulations within the spinal cord.

## Materials and Methods

### Mouse husbandry

All mice were bred and maintained in the Animal Facility of the Leibniz Institute on Aging – Fritz Lipmann Institute (FLI), Jena, Germany, according to the rules of the German Animal Welfare Law. Sex- and age-matched mice were used. Animals were housed under specific pathogen-free conditions (SPF), maintained on a 12 hour light/dark cycle, fed with mouse chow and tap water ad libitum. Mice used for analysis of fictive locomotion and projection tracing were kept according to the local guidelines of Swedish law. *Wt1^fl/fl^* mice were maintained on a mixed C57B6/J x 129/Sv strain. *Wt1-GFP* mice (Hosen et al. 2007) were maintained on a C57B6/J strain. Conditional *Wt1* knockout mice were generated by breeding *Wt1^fl/fl^* females (Gebeshuber et al. 2013) to *Nes-Cre;Wt1^fl/fl^* (Tronche et al. 1999) or *Lbx1-Cre;Wt1^fl/fl^* mice (Sieber et al. 2007). To generate mice with Wt1 ablated cells, *Wt1-GFP-DTA* mice were bred with *Lbx1-Cre* mice. Control mice were sex- and age-matched littermates (wild type or *Wt1^fl/fl^)*. For plug mating analysis, females of specific genotypes were housed with males of specific genotypes and were checked every morning for the presence of a plug. For embryo analysis, pregnant mice were sacrificed by CO2 inhalation at specific time points during embryo development and embryos were dissected. Typically, female mice between 2 and 6 months were used.

### Generation of Wt1-GFP-DTA mice

The *Wt1-GFP-DTA* mouse line bares an *IRES-lox-GFP-lox-DTA* cassette that was inserted into intron 3 of the *Wt1* locus. This cassette consists of a *GFP* encoding sequence that ends in a translational *STOP*-codon and is flanked by *loxP* sites. Downstream of *GFP*, the coding sequence for the Diphtheria toxin subunit A (DTA) was incorporated. Before Cre-induction, the IRES cassette ensures the generation of a functional GFP protein. After Cre-mediated excision of the floxed *GFP* sequence, the *DTA* is expressed from the endogenous *Wt1* promotor.

The *Wt1-GFP-DTA* model was generated by homologous recombination in embryonic stem (ES) cells. After ES cell screening using PCR and Southern Blot analyses, recombined ES cell clones were injected into C57BL/6J blastocysts. Injected blastocysts were re-implanted into OF1 pseudo-pregnant females and allowed to develop to term. The generation of F1 animals was performed by breeding of chimeras with wild type C57BL/6 mice to generate heterozygous mice carrying the *Wt1* knockin allele.

### Immunohistochemistry

Embryonic and postnatal spinal cords were dissected. They were either frozen unfixed after 15 min dehydration with 20% sucrose (in 50 % TissueTec/PBS) (post-fix) or fixed for 75 min in 4% paraformaldehyde in PBS (pre-fix). Pre-fixed tissue was cryo-protected in 10%, 20% and 30% sucrose (in PBS) before freezing in cryo-embedding medium (Neg-50 - Thermo Scientific, Kalamazoo, USA). Post- and pre-fix samples were sectioned (12 μm). Post-fixed samples were fixed for 10 min after sectioning and washed with 2% Tween in PBS (PBS-T). For pre-fixed samples, antigen retrieval was performed by incubation in sub boiling 10 mM sodium citrate buffer pH6.0 for 30 min. After blocking with 10% goat serum and 2% BSA in PBS-T (post-fix) or (prefix), sections were incubated with primary antibodies (in blocking solution) using the following dilutions: gBhlhb5 1:50 (Santa Cruz Biotechnology, Inc., Santa Cruz, California, USA), BrdU 1:100 (abcam, Cambridge, UK), shChx10 1:100 (abcam, Cambridge, UK), gpDmrt3 1:5000 (custom made (Andersson et al. 2012)), mEvx1 1:100 (1:3000 pre-fix) (Developmental Studies Hybridoma Bank, University of Iowa, Iowa City, IA), chGFP 1:1000 pre-fix (abcam, Cambridge, UK), mGFP 1:100 (Santa Cruz Biotechnology, Inc., Santa Cruz, California, USA), rFoxP2 1:800 (abcam, Cambridge, UK), mIslet1/2 1:50 (Developmental Studies Hybridoma Bank, University of Iowa, Iowa City, IA), gpLbx1 1:20,000 (gift from C. Birchmeier, MDC, Berlin, Germany), Lim1/2 1:50 (Developmental Studies Hybridoma Bank, University of Iowa, Iowa City, IA), NeuN, 1:500 (Merck, Darmstadt, Germany), rbPax2 1:50 (Thermo Fisher Scientific, Waltham, Massachusetts, USA), rbLmx1b 1:100 (gift from R. Witzgall, University, Regensburg, Germany), rbWt1 1:100 (Santa Cruz Biotechnology, Inc., Santa Cruz, California, USA). Secondary antibodies were applied according to species specificity of primary antibodies. Hoechst was used to stain nuclei. Quantitative analysis of the antibody staining was statistically analyzed using student t-test and two-way ANOVA followed by Tukey’s post hoc test.

### Bromodeoxyuridine labeling

To label proliferating cells in the embryonic spinal cord, pregnant mice at E9.5, E10.5 and E11.5 were injected intraperitoneally with 100 μg/g of Bromodeoxyuridine (BrdU) dissolved in 0.9% sodium chloride solution. Embryos were harvested at E12.5 to isolate spinal cords and stain for BrdU and Wt1. Spinal cords were frozen unfixed after 15 min dehydration with 20% sucrose (in 50 % TissueTec/PBS) and sectioned (12 μm). After any of the following treatments, sections were washed with PBS. Antigen retrieval was performed by incubation in 98°C sub boiling 10 mM sodium citrate buffer pH6.0 for 30 min. After treatment with 2N HCl at 37°C for 30 min, sections were incubated with primary antibodies using dilutions mentioned above (Immunohistochemistry). Secondary antibodies were applied according to species specificity of primary antibodies.

### RNA isolation and qRT-PCR analysis

Total RNA was isolated from E12.5 embryonic spinal cords using Trizol (Invitrogen) according to manufacturer’s protocol. Subsequently, 0.5 μg of RNA was reverse transcribed with iScript™ cDNA Synthesis Kit (Bio-Rad) and used for qRT-PCR. The primer sequences used for RT-PCR analyses are as follows: TGT TAC CAA CTG GGA CGA CA (*Act_foŕ);* GGG GTG TGG AAG GTC TCA AA (*Act_rev);* AGT TCC CCA ACC ATT CCT TC (*Wt1_qRT_for);* TTC AAG CTG GGA GGT CAT TT (*Wt1_qRT_rev)*. Real time PCR was carried out in triplicates for each sample using Syber®GreenER^TM^ (Thermo Fisher Scientific, USA) and Bio-Rad iCycler™ (Bio-Rad). PCR efficiencies of primer pairs were calculated by linear regression method. Ct values were normalized to the mean of the reference gene, *Actin*. Relative expression was determined by comparing normalized Ct values of *Wt1* conditional knockout and control samples.

### Analysis of locomotor behavior

In order to characterize gait parameters, 10 animals per sex and genotype were used. Body masses of the mice varied considerably within the groups and among the groups with significant differences between male *Wt1^fl/fl^* and *Nes-Cre;Wt1^fl/fl^* mice (*Wt1^fl/fl^:* 28 g ± 3 g vs. *Nes-Cre;Wt1^fl/fl^:* 23 g ± 3 g; Fs = 31.98; ts = 3.28, *P> 0.001*) and moderate differences between the female *Wt1^fl/fl^* and *Nes-Cre;Wt1^fl/fl^* mice (*Wt1^fl/fl^*: 25 g ± 5 g vs. *Nes-Cre;Wt1^fl/fl^:* 22 g ± 4 g; F_s_ = 3.80; t_s_ = 1.62, n.s.). We recorded the voluntary walking performance of this larger cohort using high-resolution X-ray fluoroscopy (biplanar C-arm fluoroscope Neurostar, Siemens AG, Erlangen, Germany). Because of body size variation within and among groups, we adjusted treadmill speed dynamically to the individual preferences and abilities of the mice. This method of motion analysis has been described in detail in several recent publications (e.g. Böttger et al., 2011; Andrada et al., 2015; Niederschuh et al., 2015) and will be only briefly summarized here: The X-ray system operates with high-speed cameras and a maximum spatial resolution of 1536 dpi x 1024 dpi. A frame frequency of 500 Hz was used. A normal-light camera operating at the same frequency and synchronized to the X-ray fluoroscope was used to document the entire trial from the lateral perspective. Footfall sequences and spatio-temporal gait parameters were quantified by manual tracking of the paw toe tips and two landmarks on the trunk (occipital condyles, iliosacral joint) using SimiMotion 3D.

Speed, stride length, stride frequency, as well as the durations of stance and swing phases, as well as the distances that trunk or limb covered during these phases were computed from the landmark coordinates collected at touchdown and lift-off of each limb. The phase relationships between the strides of left and right limbs as well as fore- and hindlimbs were determined from footfall sequences as expression of temporal interlimb coordination. As the animals frequently accelerated or decelerated relative to the treadmill speed, the actual animal speed was obtained by offsetting trunk movement against foot movement during the stance phase of the limb. The resulting distance was divided by the duration of the stance phase. Animal speed as well as all temporal and spatial gait parameters were then scaled to body size following the formulas published by Hof (1996): non-dimensional speed =v/gl_0_, where v is raw speed, g is gravitational acceleration and l_0_ is the cube root of body mass as characteristic linear dimension, which scales isometrically to body mass; non-dimensional frequency = f/gl0, where f is raw frequency; non-dimensional stride length = l/l_0_, where l is raw stride length. The scaled spatio-temporal gait parameters change as a function of non-dimensional speed. Therefore, linear regression analyses were computed for each parameter in the male and the female *Wt1^fl/fl^* group. The power formulas obtained from regression computation (Y = a + bX) were then used to calculate the expected value for a given non-dimensional speed for each gait parameter (baseline) in each animal of all four groups. The coefficient of determination r^2^ was computed. The deviations of the measured values of Y from the expected values, the residuals, were determined and are given in percent of deviation. Using these residuals, one-way ANOVA were computed in order to establish the significance of the differences between the means of *Wt1^fl/fl^* and *Nes-Cre; Wt1^fl/fl^* in males and females. Group means were calculated from the means of 10 animals. Sample size per mouse and limb ranged between 5 and 41 stride cycles with an average sample size of 22 ± 9.

### Fictive locomotion

Animals (P0-P3) were euthanized and the spinal cords eviscerated in ice-cold cutting solution containing (in mM): 130 K-Gluconate, 15 KCl, 0.05 EGTA, 20 HEPES, 25 Glucose, pH adjusted to 7.4 by 1M KOH) and then equilibrated in artificial cerebrospinal solution (aCSF) (Perry et al. 2015) for at least 30 minutes before the beginning of experimental procedures. Suction electrodes were attached to left and right lumbar (L) vental roots 2 and 5 (L2 and L5). A combination of NMDA (5 μM) + 5-HT (10 μM) + dopamine (DA) (50 μM) were added to the perfusing aCSF to induce stable locomotor-like output. All chemicals were obtained from Sigma. Recorded signals containing compound action potentials were amplified 10,000 times, and band-pass filtered (100-10 kHz) before being digitized (Digidata 1322A, Axon instruments) and recorded using Axoscope 10.2 (Axon Instruments Inc.) for later off-line analysis. The data was rectified and low-pass filtered using a third-order Butterworth filter with a 5 Hz cut-off frequency before further analysis. Coherence plots between L2 and L2/L5 traces were analysed using a mortlet wavelet transform in SpinalCore (Version 1.1, (Mor and Lev-Tov, 2007)). Preferential phase alignment across channels are shown in the circular plots and burst parameters were analysed for at least 20 sequential bursts, as previously described (Kiehn and Kjaerulff 1996) using an in-house designed program in Matlab (Mathworks R2014b). Ventral root recording preferential phase alignment was assessed by means of circular statistics (Rayleight test) for 20 consecutive cycles as described (Kiehn and Kjaerulff 1996). Burst parameters, including frequency, are presented as the mean ± standard deviation (SD). Burst parameters were compared using the two-tailed Mann-Whitney test or the Kruskal-Wallis analysis of variance test followed by a Dunns post-test comparing all groups.

### Tracing of commissural neurons

To examine whether the loss of *Wt1* affects spinal cord populations, tracing experiments were conducted as previously described (Rabe et al. 2009; Andersson et al. 2012). *Nes-Cre;Wt1^fl/fl^* and *Nes-Cre;Wt1++* littermate control mice P0-P5, were prepared as described above (Fictive locomotion). Two horizontal cuts (intersegmental tracing targeting commissural ascending/descending/bifurcating neurons) were made in the ventral spinal cord at lumbar (L) level 1 and between L4 and 5. Fluorescent dextran-amine (FDA, 3000 MW, Invitrogen) was applied at L1 and rhodamine-dextran amine (RDA, 3000 MW, Invitrogen) was applied between the L4/5 ventral roots. Spinal cords were incubated overnight at room temperature, subsequently fixed in 4% formaldehyde (FA) and stored in the dark at 4°C until transverse sectioning (60μm) on a vibratome (Leica, Germany).

Fluorescent images were acquired on a fluorescence microscope (Olympus BX61W1). For quantitative analyses of traced cords, consecutive images were taken between the two tracer application sites using Volocity software (Improvision, Lexington, USA). Captured images were auto-levelled using Adobe Photoshop software. Only cords with an intact midline, as assessed during imaging, were used for analysis.

Traced neurons in *Wt1^fl/fl^* control, *Nes-Cre;Wt1^fl/+^* and *Nes-Cre;Wt1^fl/fl^* cords were examined for significance using the Kruskal-Wallis analysis of variance test followed by a Dunns post-test comparing all groups. Tracing data are presented as the mean ± standard error of the mean (SEM).

### TUNEL-Assay

To detect apoptosis *in situ*, the TUNEL assay was performed prior to antibody binding. Slides were incubated with TUNEL reaction solution (1x Reaction Buffer TdT and 15 U TdT in ddH_2_O from Thermo Scientific; 1 mM dUTP-biotin from Roche) at 37 °C for 1 h and washed with PBS afterwards.

### Imaging and picture processing

Fluorescent images were viewed in a Zeiss Axio Imager and a Zeiss Observer Z1 equipped with an ApoTome slider for optical sectioning (Zeiss, Germany). Images were analyzed using the ZEISS ZEN2 image analysis software. For quantitative analyses of traced spinal cords, the application sites were identified and consecutive photographs were taken between the two application sites using the OptiGrid Grid Scan Confocal Unit (Qioptiq, Rochester, USA) and Volocity software (Improvision, Lexington, USA). Confocal images were captured on a ZEISS LSM 710 ConfoCor 3 confocal microscope and analyzed using the ZEISS ZEN2 image analysis software. Captured images were adjusted for brightness and contrast using ZEN2 image analysis software and Adobe Photoshop software.

### Statistical Analyses

Data are expressed as mean ± SD or as indicated. Groups were compared using two-way ANOVA or two-tailed two-sample equal variance student t-test as determined by group and sample size. If normal distribution of a sample was not confirmed, sample means are compared by using non-parametric Mann-Whitney *U* test. All statistical analyses were done using GraphPad Prism Software (GraphPad Software inc., San Diego, USA), IBM SPSS Statistics 24 (IMB Corporation, New York, USA), Microsoft Excel (Microsoft Corporation, Redmond, USA) or Matlab (Mathworks, R2014b). Normal distribution was assessed using the D’Agostino-Pearson normality test or Kolmogorov-Smirnov test. Significance was determined as * = P < 0.05, ** = P<0.01, *** = P<0.001.

## Acknowledgements

We thank D. Kruspe and R. Peterson for technical assistance, C. Birchmeier (MDC, Berlin, Germany) for providing the *Lbx1-Cre* mouse line and C. Hübner, H. Heuer and G. Zimmer for critical discussion. This project was supported by grants from German Federal Ministry of Education and Research (Infrafrontier grant 01KX1012) to L.B. and the Swedish Medical Research Council, Hållsten, Ländells, Swedish Brain Foundations to K.K. D.S. received a scholarship from the Leibniz Graduate School on Ageing and Age-Related Diseases (LGSA). F.V.C. was funded by a scholarship from CNPq – Brazil. The FLI is a member of the Leibniz Association and is financially supported by the Federal Government of Germany and the State of Thuringia.

## References

Andersson LS, Larhammar M, Memic F, Wootz H, Schwochow D, Rubin CJ, Patra K, Arnason T, Wellbring L, Hjalm G et al. 2012. Mutations in DMRT3 affect locomotion in horses and spinal circuit function in mice. Nature 488: 642–646.

Armstrong JF, Pritchard-Jones K, Bickmore WA, Hastie ND, Bard JB. 1993. The expression of the Wilms' tumour gene, WT1, in the developing mammalian embryo. Mechanisms of development 40: 85–97.

Burrill JD, Moran L, Goulding MD, Saueressig H. 1997. PAX2 is expressed in multiple spinal cord interneurons, including a population of EN1+ interneurons that require PAX6 for their development. Development 124: 4493–4503.

Call KM, Glaser T, Ito CY, Buckler AJ, Pelletier J, Haber DA, Rose EA, Kral A, Yeger H, Lewis WH et al. 1990. Isolation and characterization of a zinc finger polypeptide gene at the human chromosome 11 Wilms' tumor locus. Cell 60: 509–520.

Dyck J, Lanuza GM, Gosgnach S. 2012. Functional characterization of dI6 interneurons in the neonatal mouse spinal cord. Journal of neurophysiology 107: 3256–3266.

Gebeshuber CA, Kornauth C, Dong L, Sierig R, Seibler J, Reiss M, Tauber S, Bilban M, Wang S, Kain R et al. 2013. Focal segmental glomerulosclerosis is induced by microRNA-193a and its downregulation of WT1. Nat Med 19: 481–487.

Gessler M, Poustka A, Cavenee W, Neve RL, Orkin SH, Bruns GA. 1990. Homozygous deletion in Wilms tumours of a zinc-finger gene identified by chromosome jumping. Nature 343: 774–778.

Goulding M. 2009. Circuits controlling vertebrate locomotion: moving in a new direction. Nature Reviews Neuroscience 10: 507–518.

Grillner S. 1985. Neurobiological bases of rhythmic motor acts in vertebrates. Science 228: 143–149.

Grillner S, Zangger P. 1979. On the central generation of locomotion in the low spinal cat. Exp Brain Res 34: 241–261.

Gross MK, Dottori M, Goulding M. 2002. Lbx1 specifies somatosensory association interneurons in the dorsal spinal cord. Neuron 34: 535–549.

Herzer U, Crocoll A, Barton D, Howells N, Englert C. 1999. The Wilms tumor suppressor gene wt1 is required for development of the spleen. Current biology: CB 9: 837–840.

Hosen N, Shirakata T, Nishida S, Yanagihara M, Tsuboi A, Kawakami M, Oji Y, Oka Y, Okabe M, Tan B et al. 2007. The Wilms' tumor gene WT1-GFP knock-in mouse reveals the dynamic regulation of WT1 expression in normal and leukemic hematopoiesis. Leukemia 21: 1783–1791.

Kamiyama T, Kameda H, Murabe N, Fukuda S, Yoshioka N, Mizukami H, Ozawa K, Sakurai M. 2015. Corticospinal tract development and spinal cord innervation differ between cervical and lumbar targets. J Neurosci 35: 1181–1191.

Kiehn O, Kjaerulff O. 1996. Spatiotemporal characteristics of 5-HT and dopamine-induced rhythmic hindlimb activity in the in vitro neonatal rat. Journal of neurophysiology 75: 1472–1482.

Kreidberg JA, Sariola H, Loring JM, Maeda M, Pelletier J, Housman D, Jaenisch R. 1993. WT-1 is required for early kidney development. Cell 74: 679–691.

McCrea DA, Rybak IA. 2008. Organization of mammalian locomotor rhythm and pattern generation. Brain Res Rev 57: 134–146.

Moore AW, McInnes L, Kreidberg J, Hastie ND, Schedl A. 1999. YAC complementation shows a requirement for Wt1 in the development of epicardium, adrenal gland and throughout nephrogenesis. Development 126: 1845–1857.

Moran-Rivard L, Kagawa T, Saueressig H, Gross MK, Burill J, Goulding M. 2001. Evx1 is a postmitotic determinant of V0 interneuron identity in the spinal cord. Neuron 29: 385–399.

Müller T, Brohmann H, Pierani A, Heppenstall PA, Lewin GR, Jessell TM, Birchmeier C. 2002. The homeodomain factor lbx1 distinguishes two major programs of neuronal differentiation in the dorsal spinal cord. Neuron 34: 551–562.

Pearson K. 2003. Generating the walking gait: role of sensory feedback. Prog Brain Res 143: 123–132.

Perry S, Gezelius H, Larhammar M, Hilscher MM, Lamotte d'Incamps B, Leao KE, Kullander K. 2015. Firing properties of Renshaw cells defined by Chrna2 are modulated by hyperpolarizing and small conductance ion currents Ih and ISK. The European journal of neuroscience 41: 889–900.

Rabe N, Gezelius H, Vallstedt A, Memic F, Kullander K. 2009. Netrin-1-dependent spinal interneuron subtypes are required for the formation of left-right alternating locomotor circuitry. J Neurosci 29: 15642–15649.

Rackley RR, Flenniken AM, Kuriyan NP, Kessler PM, Stoler MH, Williams BR. 1993. Expression of the Wilms' tumor suppressor gene WT1 during mouse embryogenesis. Cell Growth Differ 4: 1023–1031.

Rossignol S LJ, Drew T. 1988. The Role of Sensory Inputs in Regulating Patterns of Rhythmic Movements in Higher Vertebrates - A Comparison between Locomotion, Respiration and Mastication. In: Cohen AH, Rossignol S, Grillner S, editors Neural Control of Rhythmic Movements in Vertebrates New York: John Wiley & Sons: 201–283.

Shik ML, Orlovsky GN. 1976. Neurophysiology of locomotor automatism. Physiol Rev 56: 465–501.

Sieber MA, Storm R, Martinez-de-la-Torre M, Muller T, Wende H, Reuter K, Vasyutina E, Birchmeier C. 2007. Lbx1 acts as a selector gene in the fate determination of somatosensory and viscerosensory relay neurons in the hindbrain. J Neurosci 27: 4902–4909.

Skaggs K, Martin DM, Novitch BG. 2011. Regulation of spinal interneuron development by the Olig-related protein Bhlhb5 and Notch signaling. Development 138: 3199–3211.

Tanabe Y, Jessell TM. 1996. Diversity and pattern in the developing spinal cord. Science 274: 1115–1123.

Tronche F, Kellendonk C, Kretz O, Gass P, Anlag K, Orban PC, Bock R, Klein R, Schutz G. 1999. Disruption of the glucocorticoid receptor gene in the nervous system results in reduced anxiety. Nat Genet 23: 99–103.

Wagner KD, Wagner N, Vidal VPI, Schley G, Wilhelm D, Schedl A, Englert C, Scholz H. 2002. The Wilms' tumor gene Wt1 is required for normal development of the retina. Embo J 21: 1398–1405.

Wagner N, Wagner KD, Hammes A, Kirschner KM, Vidal VP, Schedl A, Scholz H. 2005. A splice variant of the Wilms' tumour suppressor Wt1 is required for normal development of the olfactory system. Development 132: 1327–1336.

